# Directed assembly and concentrating of micro/nanoparticles, cells, and vesicles via low-power near-infrared laser generated plasmonic microbubbles

**DOI:** 10.1101/2020.01.30.927269

**Authors:** Nareg Ohannesian, Jingting Li, Ibrahim Misbah, Fusheng Zhao, Wei-Chuan Shih

**Affiliations:** Department of Electrical and Computer Engineering, University of Houston, 4800 Calhoun Road, Houston, Texas 77204, USA; Department of Biomedical Engineering, University of Houston, 4800 Calhoun Road, Houston, Texas 77204, USA; Department of Chemistry, University of Houston, 4800 Calhoun Road, Houston, Texas 77204, USA; Program of Materials Science and Engineering, University of Houston, 4800 Calhoun Road, Houston, Texas 77204, USA

**Keywords:** Microbubbles, Plasmonic Nano-heaters, Particle Manipulation, Selective Particle Assembly, Extracellular vesicles

## Abstract

Directed assembly and concentrating of micro- and nanoparticles via laser generated plasmonic microbubbles in a liquid environment is an emerging technology. For effective heating, visible light has been primarily employed in existing demonstrations. In this paper, we demonstrate a new plasmonic platform based on nanoporous gold disk (NPGD) array. Thanks to the highly tunable localized surface plasmon resonance of the NPGD array, microbubble of controlled size can be generated by near-infrared (NIR) light. Using NIR light provides several key advantages over visible light in less interference with standard microscopy and fluorescence imaging, preventing fluorescence photobleaching, less susceptible to absorption and scattering in turbid biological media, and much reduced photochemistry, phototoxicity and whatsoever. The large surface-to-volume ratio of NPGD further facilitates the heat transfer from these gold nanoheaters to the surroundings, achieving unprecedented low-power operation. While the microbubble is formed, the surrounding liquid circulates and direct microparticles randomly dispersed in the liquid to the bottom NPGD surface, yielding unique assemblies of microstructures. Such capability can also be employed in concentrating suspended colloidal nanoparticles at desirable sites and with preferred configuration, both enhancing the sensor performance. In addition to various micro- and nanoparticles, the plasmonic microbubbles are also shown to collect biological cells and nanovesicles. By using a spatial light modulator (SLM) to project the laser in arbitrary patterns, parallel assembly can be achieved to fabricate an array of clusters. These assemblies have been characterized using optical microscopy, scanning electron microscope, hyperspectral localized surface plasmon resonance imaging and hyperspectral Raman imaging.

## Introduction

The ability to remotely manipulate micro- and nanoparticles has been a tantalizing capability. Many techniques have been demonstrated using forces and fields of various origins. Electrokinetic mechanisms including electrophoresis (EP), dielectrophoresis (DEP), and electroosmotic flow have been employed to separate and/or concentrate cells according to their bulk and/or surface electrical properties [1]. Acoustic mechanisms such as standing bulk/surface wave and traveling wave have also been implemented [2]. When cells are tagged with magnetic materials, they become manipulatable by magnetic fields [3]. Optical tweezers use gradient force to manipulate particles [4]. Optically induced dielectrophoresis (ODEP) further combines light with dielectrophoresis [5]. Recently, several studies have utilized photothermally generated microbubbles for particle manipulation. Microbubbles can be generated upon laser irradiation of metal films and the Marangoni flow around the bubble can drive particles to the 3-phase interface [6–8]. Microbubble initiation occurs through a mechanism called explosive expansion of superheated liquids where vaporization and dissolved gas in the medium combine and form the bubble [9]. Degassed water is still capable of photothermal microbubble generation, however the overall saturated bubble is reduced in size [10]. Using a plasmonic substrate as the light absorber to generate microbubbles has considerably reduced the necessary laser power to generate a plasmonic microbubble and successfully assembled micro- and nanoparticles for on-chip sensing devices [11].

Localized surface plasmon resonance (LSPR) is the light-excited collective oscillation of conduction band electrons confined within coinage metallic nanostructure. LSPR leads to localized and enhanced electric fields near the nanostructure surface. Such electric field localization has been widely recognized as the primary mechanism in surface-enhanced optical phenomena involving one photon such as surface-enhanced infrared absorption (SEIRA) [12], surface-enhanced near-infrared absorption (SENIRA) [13], and two photons such as surface-enhanced Raman scattering (SERS) [14] or metal-enhanced fluorescence (MEF) [15]. The localization of highly enhanced electric fields, also known as ‘‘hotspots”, has profound impact to light-matter interactions on plasmonic nanostructures. Nano-gaps of less than 10 nm in nanostructures or between nanoparticles support gap-mode LSPR and induce remarkable electric field localization [16, 17]. Various methods have been proposed and implemented for nano-gap generation via lithography patterning of nano-gaps in gold film [18], nano-gap formation between nanoparticle or nanowire assembly [19, 20], gold nanoparticle (AuNP) aggregation [21], and bi-continuous gold ligaments and porous network in nanoporous gold [22].

Most studies reported on photothermal or plasmonic microbubble generation either utilizes a thin layer of gold film [7, 8, 23, 24] or gold nanoparticles [9, 25, 26] with visible laser sources (532 nm) as the primary choice to maximize absorption. There are a few exceptions where a near-infrared laser source was used [27, 28]. Tantussi et al. generate and guide a microbubble on a nanoantenna using a femtosecond laser (30 to 150 mW). In contrast to using a gold film, the microbubble can only be generated on where the nanoantenna is located. The packing density of nanostructures will also influence the efficiency of transferring laser power to heat. Regardless of particles size, high power lasers are necessary for photothermal and plasmonic microbubble generation [7, 9, 24–27, 29].

Our interest lies in the plasmonic microbubble generation on nanoporous gold disk (NPGD) array with unprecedented low-power operation (5-30 mW). NPGD features unique structural hierarchy, consisting of tunable external geometry and internal bi-continuous ligaments and pores [30]. The semirandom porous network has a characteristic length of 5-15 nm and has been shown to provide significant plasmonic field localization. NPGD array has been demonstrated to be highly effective in surface-enhanced Raman spectroscopy [31], surface-enhanced fluorescence [32], surface-enhanced near infrared absorption [13], plasmon-enhanced heterogeneous catalysis [33], and photothermal heating [10, 34, 35]. Thanks to the highly tunable LSPR of the NPGD array, microbubble of controlled size can be generated by near-infrared (NIR) light. Using NIR light provides several key advantages for bio applications over visible light in less interference with standard microscopy and fluorescence imaging, preventing fluorescence photobleaching, less susceptible to absorption and scattering in turbid biological media, and much reduced photochemistry, phototoxicity and whatsoever [36]. Recently, we reported the generation of microbubbles on NPGD using 785 nm laser output with holographically generated patterns. We investigated the dynamics of microbubble formation and disappearance with respect to laser power and degassing conditions [10]. However, the onset of the microbubble formation was not available due to imaging speed limit. In this paper, we first demonstrate an improved imaging system to better capture the microbubble formation transient. We further investigate how the microbubble and its surrounding flow would interact with particles of different size and surface properties. For potential applications, we demonstrate the feasibility of forming unique microparticle assemblies, separating hydrophilic and hydrophobic particles, concentrating gold nanoparticles, biological cells and vesicles at prescribed locations for improved sensing. We further show the feasibility of providing patterned illumination by a spatial light modulator for parallel operations of forming an array of assemblies.

## Materials and Methods

### Materials

Sodium citrate dehydrate (≥99.0%), gold(III) chloride hydrate (99.999% trace metals basis), sodium dodecyl sulfate (ACS reagent, ≥99.0%), 40 nm AuNPs, latex beads (polystyrene beads), 10% aqueous suspension) with mean particle sizes of 300 nm and 750 nm, and 1,2-Di(4-pyridyl)ethylene (BPE, 97%) were purchased from Sigma-Aldrich. Ethanol (200 proof) was purchased from Decon Laboratories, Inc. Ag70Au30 (atomic percentage) alloy sputtering target was purchased from ACI Alloys, Inc. Argon gas (99.999%) was used for RF-sputter etching.

### Methods

#### Fabrication of NPGD substrate

First, a 2 nm titanium film and a 3 nm gold film were evaporated onto a clean glass coverslip as an adhesion layer. An 80 nm thick Au-Ag alloy film was then sputter deposited onto the coverslip using an alloy target (Au30%Ag70%, ACI Alloy). A single layer of 300 nm diameter polystyrene (PS) beads (Sigma-Aldrich) was formed on top of the alloy film, followed by a timed oxygen plasma treatment to shrink the PS beads and guarantee the separation of neighboring beads. The bead pattern was sputter etched onto the alloy film using argon plasma. Once the pattern transfer was complete, the PS beads were removed by sonication in deionized (DI) water. The samples were dealloyed in 70% nitric acid for 1 minute, followed by a 2 min DI water rinse. The fabrication conditions produce as-prepared NPGDs with 230 nm diameter, 80 nm thickness and 5-15 nm pore size. A more detailed fabrication process of NPGDs was described in our previous work [22].

#### Preparation of PS beads and AuNPs

Both PS beads (Sigma-Aldrich) and AuNPs (Sigma-Aldrich) had a hydrophilic surfactant layer to be stored as stable suspension. To remove the surfactant for PS beads, the solution was washed as follows: centrifugation at 3000 RPM for 20 min; removal of clear top layer of solution; redispersion in DI water. The washing steps were repeated three times to make the PS beads hydrophobic.

#### Optical setup

A 7 mm beam from a 785 nm continuous wave (CW) laser (Spectra Physics 3900S) was sent to a SLM (Boulder Nonlinear XY Phase Series) that performed phase modulation and produce the desired pattern on the sample plane of an inverted microscope (Olympus IX71). The phase holograms were pre-calculated and loaded into the SLM. The modulated beam cast an illumination pattern onto the sample through the back port and the objective lens (60 X water immersion) of the microscope. The translucency of the NPGD substrate enables simultaneous imaging and heating from either above or below the substrate. The transmitted light was then sent through the side port of the microscope to an electron multiplied charge coupled device (EMCCD) camera (ProEM, Princeton Instruments) for bright field imaging and high speed recording at 100 fps [37]. The SERS map of BPE was produced using a home-built line-scan Raman microscope [38]. The recorded data were subjected to an automated image curvature correction algorithm, followed by 5^th^-order polynomial background removal [39]. A home-built hyperspectral extinction microscope system, consisting of an inverted microscope (Olympus IX71) and a spectrograph (Acton SP2300, Princeton Instruments) mounted with a CCD camera (PIXIS 400BR, Princeton Instruments), was used to acquire an extinction spectra map of the AuNP modified NPGD array on a glass coverslip. A broad band halogen lamp and a bright-field condenser were used to illuminate the sample. The transmitted image of the sample was projected through a 40 X objective lens onto the 20 μm entrance slit of the spectrograph. The transmitted image was laterally scanned for 2D imaging with 20 μm step size, resulting in an extinction map with 0.5 × 0.5 μm^2^ spatial resolution.

#### Laser power control

To precisely control the laser, the SLM was used as a frequency-agile power modulator by controlling its on/off duty cycle. The SLM (maximum modulation frequency at 333 Hz) was set to display two patterns alternatively with one irradiated at the region of interest while the other outside the EMCCDs imaging range. The lowest possible setting for plasmonic microbubble production was achieved by setting a duty cycle of 1/10 which reduced the average laser power from 60 mW/spot to a nominal 6 mW/spot. To determine a threshold power to generate identical microbubbles at every location on the NPGD array, the duty cycle was slowly increased while monitoring the laser spot on the sample plane. At 8 mW/spot the bubble exceeded the desired diameter of 25 μm on the sample. Henceforth the duty cycle of the SLM was fixed to maintain an 8 mW/spot laser power.

#### *E. coli* and Extraction of exosomes

*E. coli* was purchased from ATCC and was cultured in LB Broth for 12 hours at 37 ^0^C. Exosomes were collected from 5 mL of media from lung cancer cell culture (2-4× 10^6^ cell/ml, cell line H460). The culture media were collected, subjected to centrifugation at 800 × g for 10 min to sediment the cells, and centrifuged at 12,000 × g for 45 min to remove the cellular debris. The exosomes were separated from the supernatant via centrifugation at 100,000 × g for 2 h [40]. The exosome pellet was washed once in a large volume of PBS and resuspended in 100 μL of PBS to yield the exosome fraction. The concentration of extracted exosomes was quantified using NanoSight particle tracking (Malvern).

## Results and discussion

### Mechanism for particle assembly

All assembled particles, AuNPs (40 nm) and PS beads (750 nm), retained their hydrophilic surface to mimic biological samples. A separate batch of PS beads (750 nm) were treated to present a hydrophobic surface as described in the Materials and Methods. The NPGD substrate was immersed in the colloidal solution containing the particles, sandwiched between two No. 1 glass coverslips and placed facing down on the water immersion objective lens (60X) of the microscope (Fig. 1a-b). The gap between the two coverslips was 100 μm as maintained by a spacer. Since hydrophilic particles remained suspended in the medium while hydrophobic particles sunk in the solution, having the NPGD substrate facing down can effectively prevent unwanted particle settlement on the substrate.

**Fig. 1.**
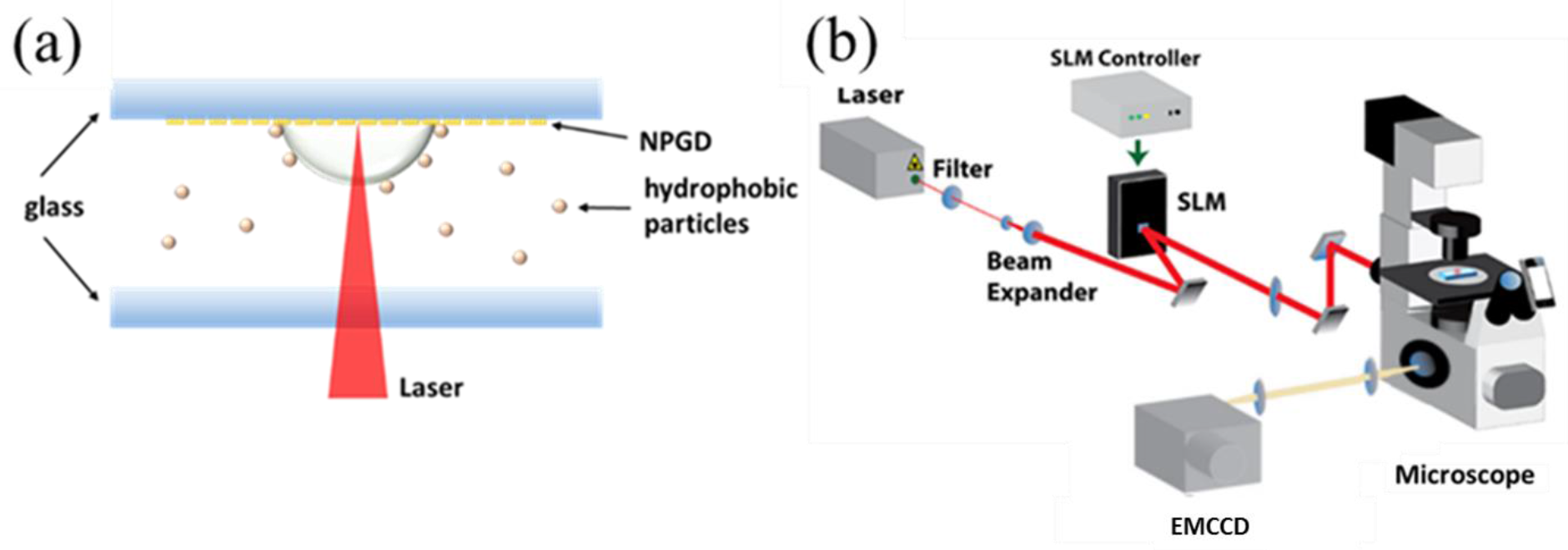
(a) Sample setup. The NPGD are fabricated on a coverslip glass that is facing down on an inverted microscope; (b) Schematic of optical setup.

The plasmonic microbubble assembly works as follows: The laser light (785 nm) was focused onto the NPGD substrate with a diameter of 1.5 μm. NPGDs can then convert the laser power into heat plasmonically due to a higher thermal efficiency as demonstrated previously. The microbubble formed immediately upon laser exposure (8 mW) and expanded to 15 μm in diameter within a sec after which it continued to increase in size at a reduced rate (Fig. 2). Upon reaching a steady state, chemical potential of dissolved gas is achieved preventing any further permeation of gasses at the microbubble interface. An imaging speed of 100 fps was used to capture the onset of microbubble formation (<10 μm), which was not possible previously (Fig. 2) [10]. The laser was set to focus on the surface of the NPGD array in order to achieve maximum absorption for microbubble generation, while a 4f imaging system was used to image roughly 5 μm above the surface to clearly image the microbubble growth dynamics. In the presence of the substrate, the microbubble has a three quarter circle bound to the plasmonic substrate [9]. Microbubbles first appeared as a hazy circle of about 2.5 μm. After 0.01 sec, the combination of dissolved gas/water vapor causes continued volume expansion, reaching 12 μm in size after 0.8 sec. Power law (*14.12*t^0.14^*) was used to fit the diameter vs. time as shown in Fig. 2.

**Fig. 2.**
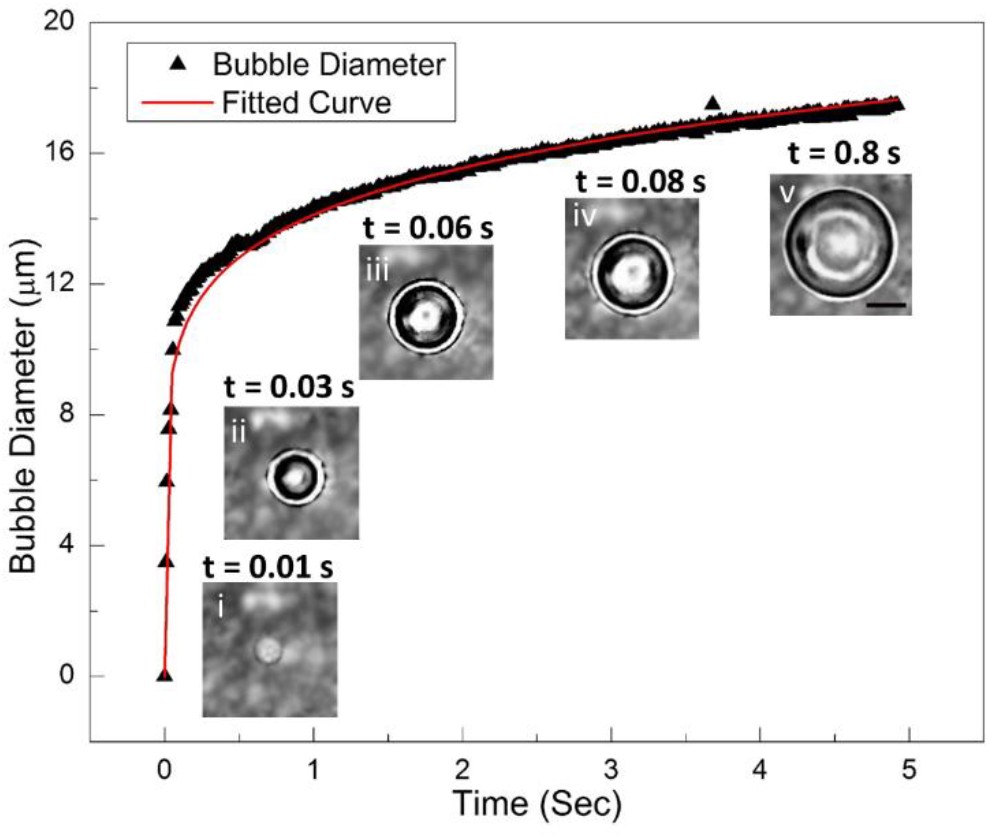
Microbubble growth curve vs time with power law fit with Consecutive bright-field images of microbubble formation under 8 mW laser power.

A convective flow in the surrounding liquid directs the particles to the interface between the plasmonic microbubble and the bottom substrate. The primary mechanisms contributing to generating a convective flow are temperature gradient and surface tension induced Marangoni convection, both of which are responsible of particle collection and entrapment at the boundaries of the bubble [23]. When the laser was turned off, the microbubble shrank and eventually disappeared as characterized previously. Some particles adhered to the gas-liquid interface were eventually gathered on the substrate by the shrinking microbubble.

### Plasmonic microbubble enabled assembly of 750 nm polystyrene beads

Particle assembly was characterized using both PS beads and AuNPs at different bubble sizes. The bubble on the NPGD substrate was maintained for roughly 30 sec, which provided sufficient time to reach equilibrium for particle entrapment. After particle assembly, the laser spot was removed from the substrate, allowing the bubble to shrink and disappear and leaving the particles fixed onto the substrate. The substrate was then gently washed with DI water and dried using nitrogen gas. Particle assemblies were then imaged by SEM.

Previous publications on microbubble assembly mainly focus on inorganic particles such as quantum dots and silica beads which are mainly hydrophobic [23]. However, since most biological cells and vesicles are hydrophilic, we investigated the impact of surface tension with the medium on overall particle collection. PS beads were washed using steps listed in the materials and methods section. Removing the surfactant layer from the surface restores the hydrophobic nature of the PS beads. A plasmonic microbubble (25 μm) was generated in two separate media, one containing hydrophobic particles and the other containing hydrophilic particles. Hydrophilic particles were immediately collected upon plasmonic microbubble initiation. Upon laser cutoff and the complete shrinkage of the microbubble revealed a cluster of PS beads as seen in Fig. 3a. Hydrophobic PS beads underwent a similar collection process upon plasmonic microbubble resulting in the cluster seen in Fig. 3b. Removing the surfactant layer showed no difference in particle assembly, suggesting that biological samples can be manipulated by plasmonic microbubbles (Fig. 3a-b).

**Fig. 3.**
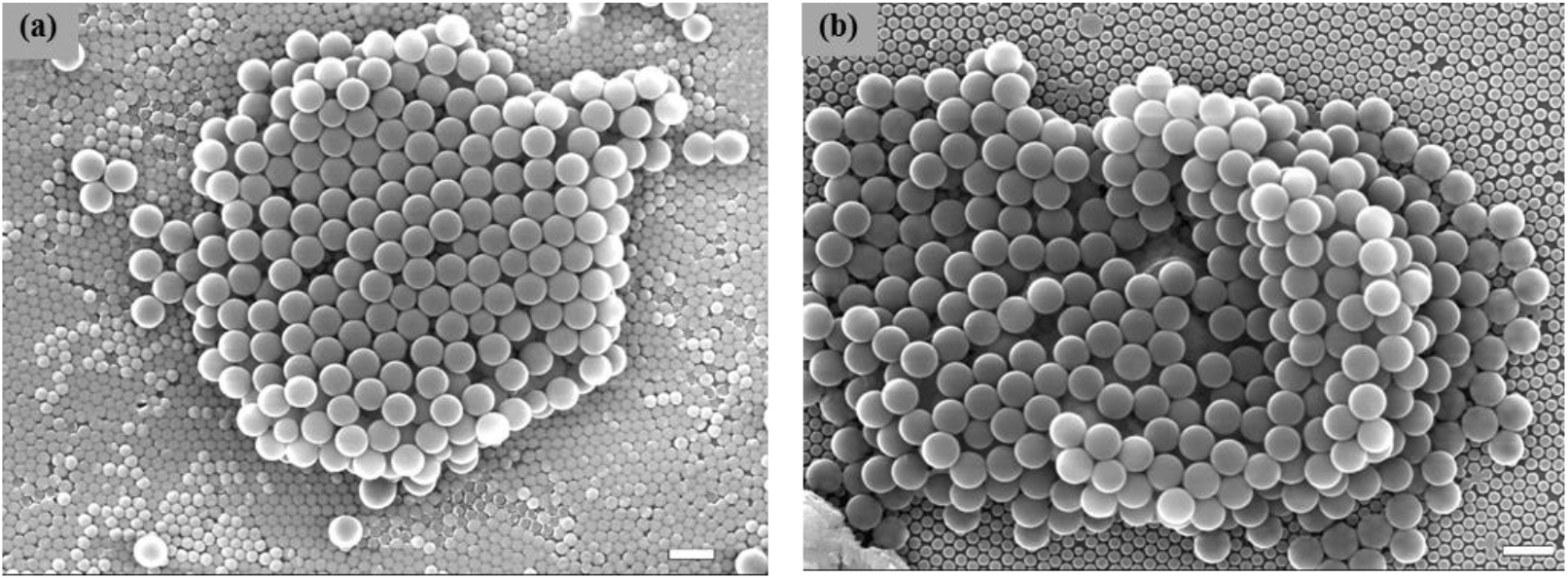
SEM images of (a) hydrophilic and (b) hydrophobic surface PS beads collected by plasmonic microbubble. Scale bar: 1 μm.

Further investigation was done on the effect of bubble size on particle assembly. Microbubbles with diameters ranging from 3 μm to 20 μm were generated (Fig. 4). Bubble diameters smaller than ~5 μm did not form a well-defined assembly shape due to the comparable length scale between the bubble and the PS beads. The PS beads trapped on the bubble do not have enough space to reorganize themselves into a closely packed arrangement. Removing the microbubble resulted in random PS beads stacks in irregular 3D structures (Fig. 4a). For bubbles with diameters around 5 μm, trapped PS beads organized themselves into closely packed hexagonal patterns. After the removal of the microbubble, the PS beads formed a 3D hollow dome structure. The 3D structure’s hollow state is strongly indicated through a lack of clear layer structure, instead of the multilayer stacking as observed from the top-view SEM image (Fig. 4b) and the 45° angle view (Fig. 4g). For bubble sizes between 5 μm and 10 μm in diameter, the small size of the PS beads caused the dome structure to collapse once the bubble was removed (Fig. 4c, h). As bubble diameter surpasses 10 μm, the assembled particles correspondingly reveal larger caved in regions with increased bubble size (Fig. 4d, i). For bubbles with diameter of 20 μm or larger, the PS beads are too small to support any hollow 3D structure which results in single or multilayer flattened structures (Fig. 4e, j). The dome structure appears regardless of hydrophilic or hydrophobic state (Fig. 4f).

**Fig. 4.**
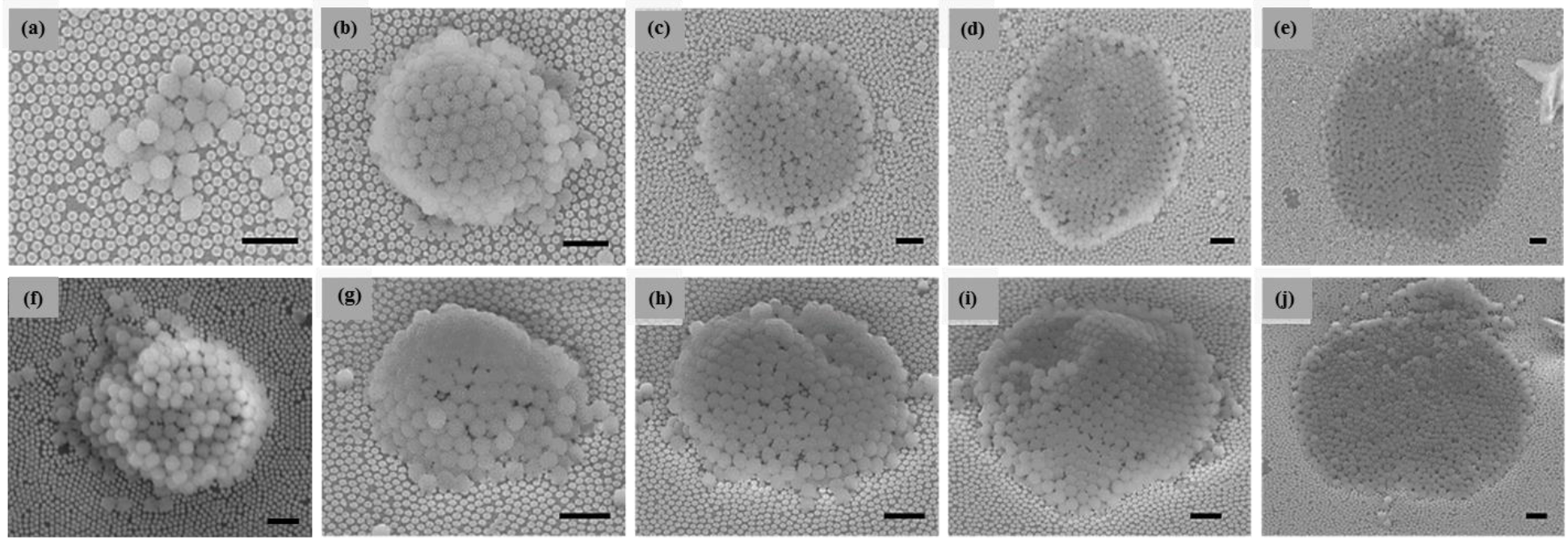
Effect of plasmonic microbubble diameter on PS beads assembly. (a-e) top view of hydrophobic surface PS beads assembly; (f) top view of hydrophilic surface PS beads assembly; (g-j) 45° view of hydrophobic surface PS beads assembly shown in (b-e) respectively. Scale bar: 2 μm.

### Variation in particle assembly regarding hydrophilic and hydrophobic surfaces

As mentioned earlier, a plasmonic microbubble forms a convective flow in its near vicinity that transports water and dissolved air molecules towards the laser focal point on the substrate. Since hydrophilic particles have higher affinity to water molecules. We hypothesize that the water molecules act as carriers transporting any colloidal particles towards the microbubble. In contrast, hydrophobic particles tend to avoid water molecules. They adhere to the substrate or dissolved gas molecules in the medium. Thus, hydrophobic particles tend to use dissolved air molecules as transport carriers towards the microbubble. To test this hypothesis, we performed experiments of particle assembly of 750 nm PS beads after degassing the water and compared with previous results without degassing. We found no significant change in hydrophilic particles assembly in the degassed experiment (Fig. 5). However, hydrophobic particles did not form any assembly in the degassed experiment, suggesting that the presence of gas molecules are essential in the assembly process for hydrophobic particles.

**Fig. 5.**
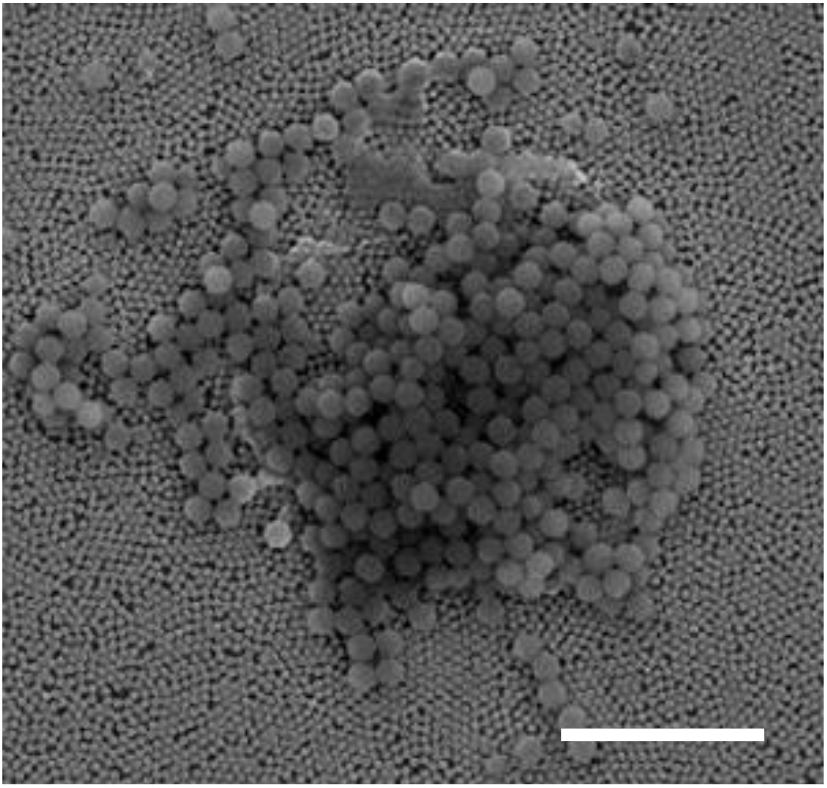
Plasmonic microbubble enabled collection of hydrophilic surface PS beads in degassed water. Scale bar: 5 μm.

### Selected surface modification using 40 nm AuNPs

After assembled on the NPGD array, the polystyrene beads cause a local index change that can alter the LSPR of NPGD underneath. It is also known that metal nanoparticles can induce plasmonic coupling due to nanogaps generated from close proximity to the NPGD array. Plasmonic coupling alters the extinction properties and can generate stronger electric fields to be used for various application such as surface-enhanced Raman scattering (SERS) and other techniques [30]. To demonstrate the plasmonic interactions among assembled particles and with the underlying NPGD array, hydrophobic 40 nm gold nanoparticles were deposited on the NPGD array using different microbubble sizes to monitor cluster formation. No hollow 3D structures were formed with AuNPs regardless of bubble size, presumably due to the significant size mismatch between the microbubble and the AuNPs. Similar to the PS beads, particle assembly was unaffected by the hydrophobic or hydrophilic state of the AuNP without degassing (Fig. 6). The most common form of AuNP particle assembly was the ring structure. Microbubbles smaller than 5 μm in size may result in deformed ring structures (Fig. 6a-b). On the contrary, microbubbles greater than 5 μm formed well-defined ring structures, which generally scales with the microbubble diameter. Larger microbubbles displayed a phenomena similar to the “coffee ring” effect which deposits the majority of AuNPs at the edge of the microbubble (Fig. 6c-d) [41].

**Fig. 6.**
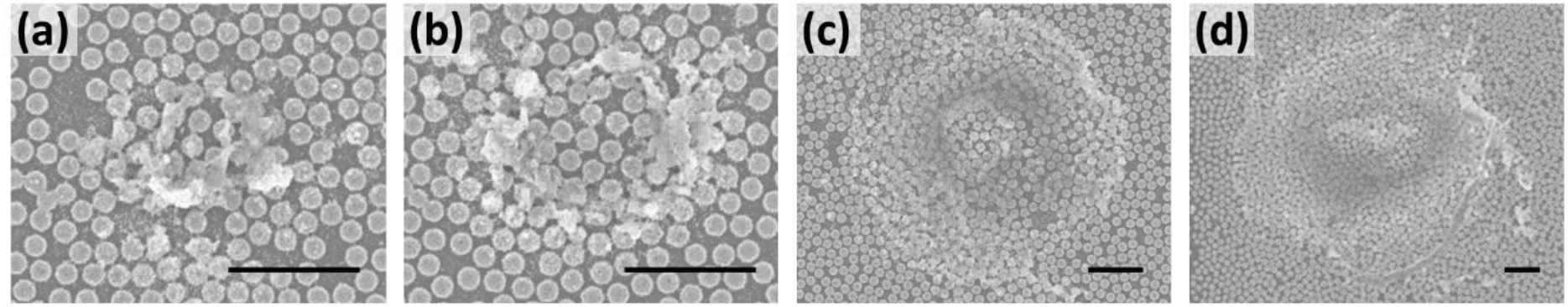
Plasmonic Microbubble enabled gold nanoparticles assembly. Scale bar: 2 μm.

Beyond providing an accurate power control mechanism, the SLM also provides the capability of multiple laser spots at precisely controlled location and real-time reconfigurability. A 4 × 4 AuNP cluster array was formed onto the NPGD substrate by a single SLM pattern without any movement of the sample (Fig. 7a). In other words, an array of 16 microbubbles of 5 μm were generated simultaneously for parallel operation.

**Fig. 7.**
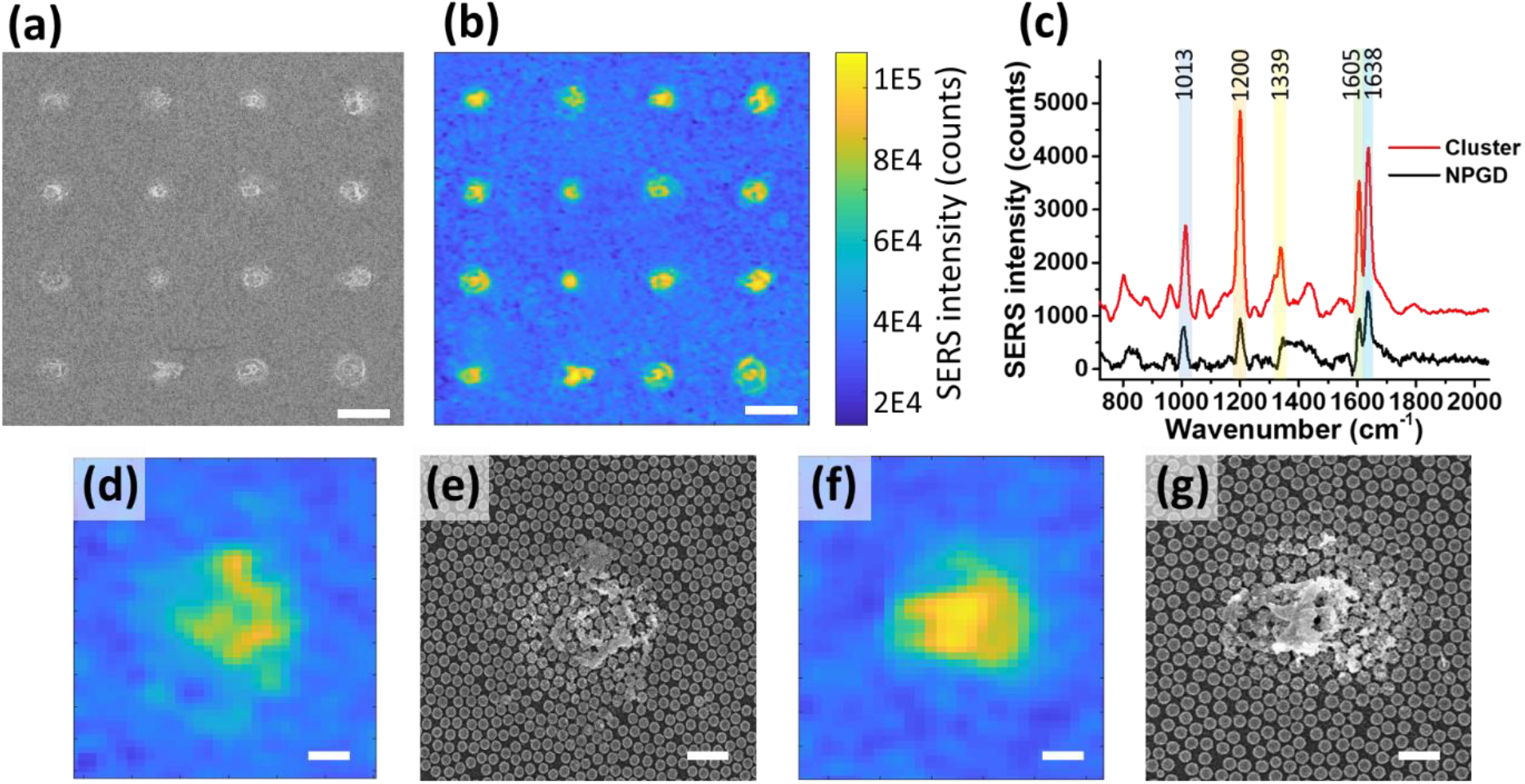
(a) SEM of 4 by 4 AuNP cluster array on NPGD substrate. (b) Corresponding SERS map of BPE plotted at 1200 cm^-1^ peak for (a). (c) SERS intensity of BPE on AuNP cluster and on NPGD. (d, f SERS map of single AuNP cluster. (e, g) SEM image of (e,f). Scale bar: (a-b) 10 μm, (d-g) 1 μm.

### SERS map study

Aggregated metal nanoparticles form strong electric field hot-spots due to nanogap coupling effects, which has been utilized for SERS measurements. Although traditional aggregation relies on chemical precipitation, here we demonstrate the plasmonic microbubble approach. The SERS measurement was carried out using the 4 × 4 AuNP cluster array shown previously (Fig. 7a). The sample was incubated in 1 mM BPE solution for 1 hour, afterwards thoroughly washed with 200 proof ethanol and blow dried by nitrogen gas flow. The SERS map was obtained using a home-built line-scan Raman system as described in the Materials and Methods section. SERS spectra of BPE obtained from bare NPGD substrate and AuNP clusters reveal prominent peaks at frequencies 1013, 1200, 1339, 1605, and 1640 cm^-1^ (Fig. 7c). The area under the curve of the 1200 cm^-1^ peak was used to generate the SERS map that shows good agreement with the SEM images (Fig. 7 a-b, d-g). The map shows AuNP clusters with approximately 5 times the SERS intensity of bare NPGD (Fig. 7c), which has been shown to have a SERS enhancement factor exceeding 10^8^ [31]. The outer region of AuNP clusters appear to have a thin layer of AuNP attached to the NPGD that shows higher SERS signal compared to bare NPGD (Fig. 7 d-e). The enhanced SERS activity indicates an increase in hot-spots on the surface induced by a strong electromagnetic coupling effect between AuNPs and the NPGD array. Regions with larger AuNPs aggregation triggers an additional gap-mode coupling between AuNPs alongside AuNPs-NPGD coupling that further enhances the SERS signal (Fig. 7 f-g). The gap-mode coupling is due to Nano-gaps between closely packed nanoparticles which does not exist when particles are scattered or in a colloidal suspension. Increased SERS signal can be expected as cluster size increments but sizes above 200 nm in thickness has limited coupling with the surface beyond which the SERS intensity plateaus.

### Hyperspectral extinction peak map study

To further study the coupling effect between the assembled AuNPs on NPGDs, we acquired an SEM image and its corresponding extinction peak map over the same AuNP cluster as shown in Fig. 8. The extinction map was obtained using the home-built instrument described in the Materials and Methods section. First, by comparing Fig. 8 a and b, we observe that the general structural architecture of the cluster match quite well in both images. Significant extinction peak redshift has been observed from corresponding locations where AuNPs are identified in the SEM image. We further examine three structurally distinct regions in these images. Region I was the bare NPGDs as shown in Fig. 8d with an average extinction spectrum shown in Fig. 8c (“i” in green) peaking at ~620 nm. The peak position of the NPGD substrate was governed by both nanodisk diameter and particle packing density [42]. The extinction spectra in Fig. 8c were normalized to its maximum intensity. Region II is where the NPGDs were covered by a thin layer of AuNPs and strong plasmonic coupling occurs between AuNPs and NPGDs (Fig. 8e). Compared to bare NPGDs, the extinction spectrum of this region is redshifted by ~50 nm as shown in Fig. 8c (“ii” in black). Region III is where a thick layer of AuNP cluster covers the NPGDs (Fig. 8f). Due to the significant clustering, the extinction spectra shown in Fig. 8c (“iii” in red) is dominated by AuNP-AuNP and AuNP-NPGD coupling which resulted in an average redshift of ~100 nm compared to bare NPGDs. The red shift is a strong evidence of near-field coupling effect among AuNPs as well as between AuNPs and NPGDs, which provides additional localized electric field enhancement for the highest SERS signal discussed previously in Fig. 7 [35]. An extinction spectrum of colloidal (non-clustered) AuNPs reveals a peak at ~520 nm indicating the lack of AuNP-AuNP plasmonic coupling without clustering (Fig. 8c “iv”). In other words, the clustered AuNPs on NPGDs in Region III exhibited a redshift of 200 nm compared to colloidal, nonclustered single AuNPs.

**Fig. 8.**
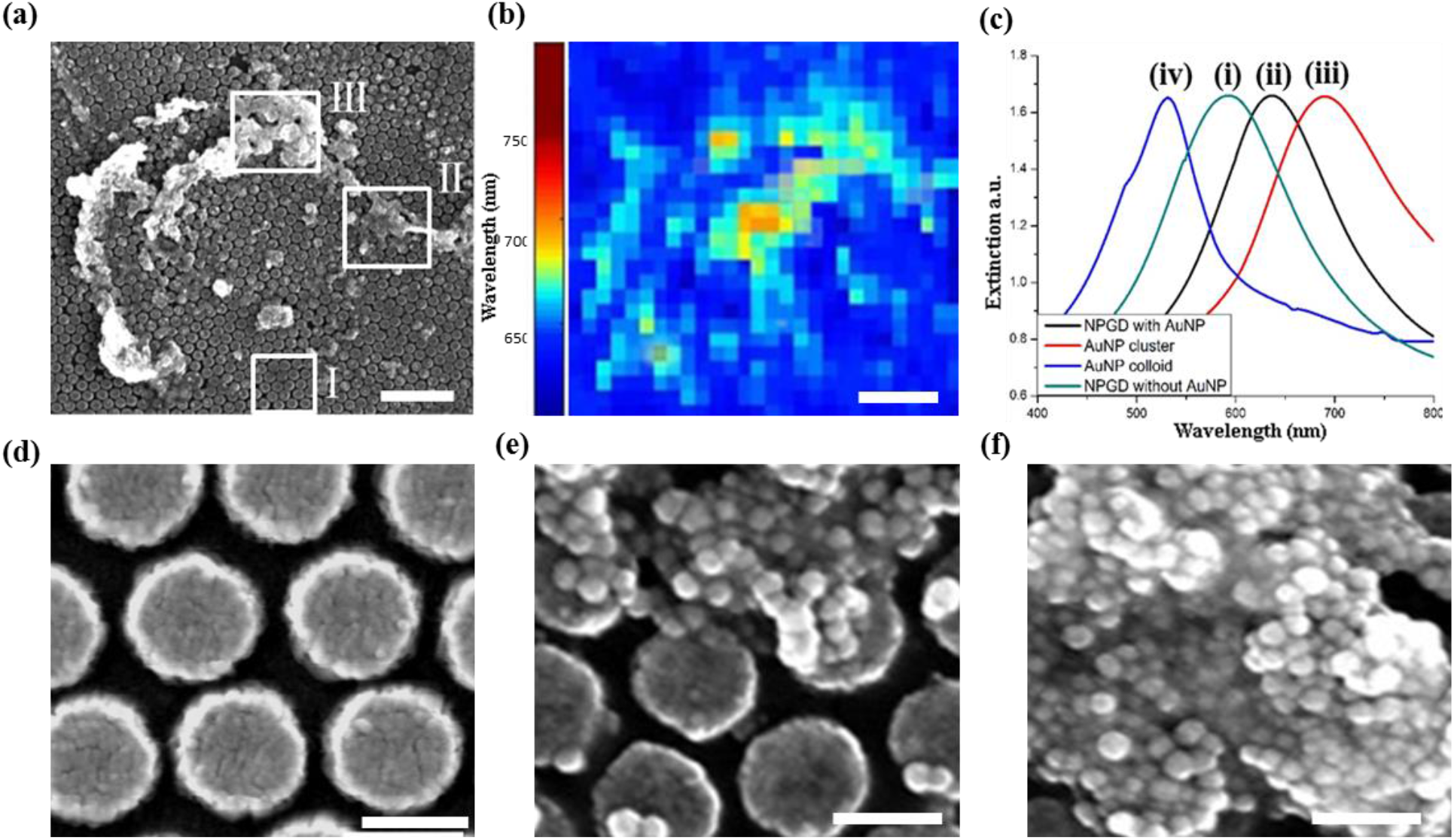
(a) SEM image of AuNP cluster; (b) Corresponding extinction peak map of (a); (c) Average extinction spectra obtained from various regions and colloidal AuNPs. Zoomed-in SEM images of (a) showing (d) region I; (e) region II; (f) region III. Scale bar: (a-b) 1 μm;(d-f) 200 nm.

### Collection of biological organisms

To demonstrate the ability of plasmonic microbubble enabled assembly and concentration of biological cells and vesicles, *E. coli* suspended in aqueous solution was drop cast onto the NPGD array (Fig. 9a). Generation of plasmonic microbubble triggered the immediate collection of *E. coli.* The complete shrinkage of the microbubble revealed a cluster of *E. coli* indicating a successful collection of living organisms with no sign of damage (Fig. 9b).

**Fig. 9.**
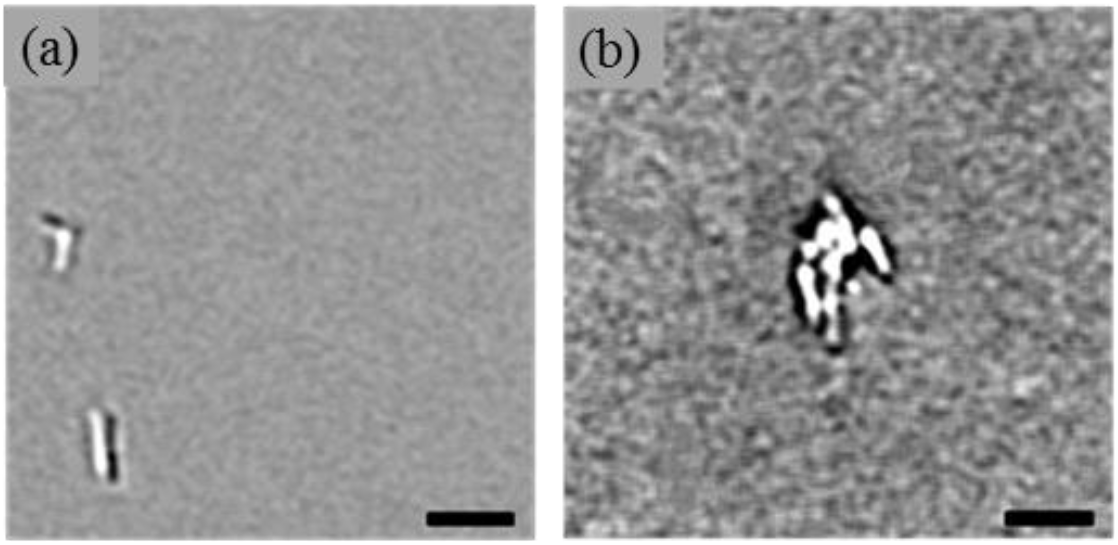
Bright-field images of E. coli collection in LB Broth; (a) E. coli roaming around with Brownian motion before plasmonic microbubble generation; (b) Cluster of E. coli after plasmonic microbubble completely gone. Scale bar: 5 μm.

Upon the successful collection of *E. coli*, which ranges between 5 μm to 10 μm, we divert our attention toward collection of biological organisms that are in nanoscale. Exosomes are extracellular vesicles known for carrying abundant dsDNA, mRNA and miRNA of the source cells. Their size ranges from 50 to 250 nm and are excreted by every mammalian cell. Recent evidence suggests that dysregulation of the genetic content within exosomes has a major role in tumor progression and in the surrounding microenvironment [43]. However, the nanoscale size of exosomes renders it to remain in a colloidal suspension which makes effective capture and concentration challenging. Herein, we demonstrate the capability of plasmonic microbubbles for aggregating and concentrating exosomes. Single nanoparticle tracking (NanoSight, Malvern) has been employed to produce a size histogram as shown in Fig. 10a, where the average size was 151 nm with a standard deviation of 61 nm. A plasmonic microbubble of 25 μm was generated and left for 2 min (Fig. 10 b-c). After the complete shrinkage of the microbubble, a large quantity of exosomes was accumulated indicating a successful aggregation and concentration process (Fig. 10d). It is worth noting that the collected exosomes appeared to be surrounding the microbubble, but not within it.

**Fig. 10:**
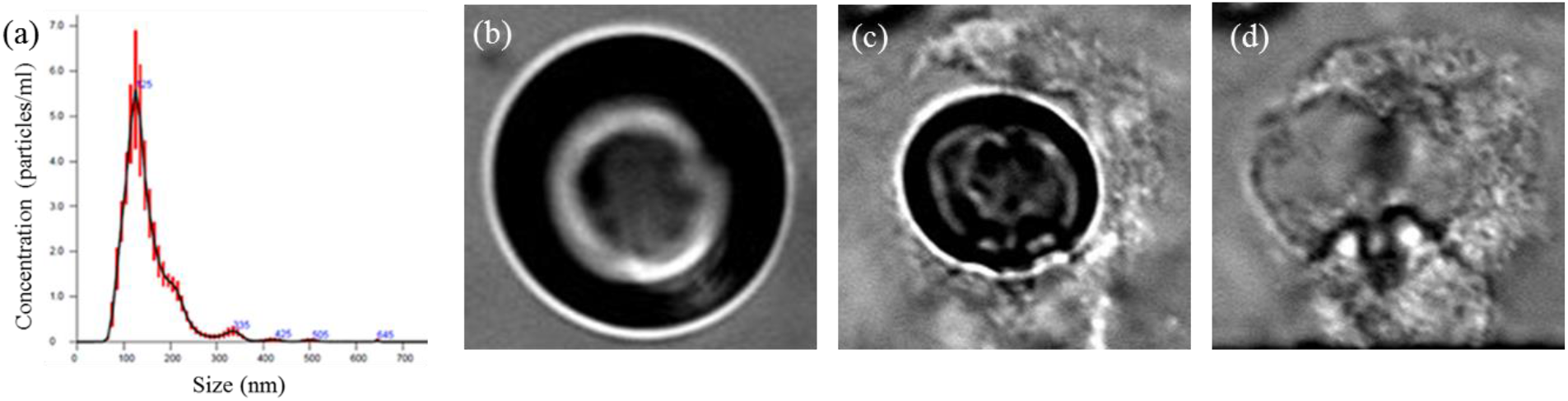
(a) Size distribution of exosome determined using NanoSight nanoparticle tracking analysis. Bright-field images of exosome collection in PBS 1X buffer; (b, c) Generated plasmonic microbubble for exosome collection left for 10 sec and 120 sec respectively; (d) Cluster of collected exosome after shrinking the microbubble completely. Scale bar: (a-c) 2 μm.

## Conclusion

In summary, we have presented a novel platform for directed assembly and concentrating of micro/nanoparticles in solutions via low-power near-infrared (NIR) laser generated plasmonic microbubbles on nanoporous gold disk array. NIR illumination provides several advantages over visible illumination in lower phototoxicity, less interference to fluorescence imaging, and more immune to solution turbidity. The technique showed no limitation upon changing from hydrophobic to hydrophilic particles. We hypothesized and confirmed that hydrophilic particles rely on water molecules as carrier towards the microbubble while hydrophilic particles relied on dissolved air molecules, indicating that surface tension plays an important role on particle transport mechanism. Based on this principle, it is possible to selectively collect hydrophilic particles by excluding hydrophobic particle collection through degassing of the medium. Upon varying the microbubble size, different forms of clusters formation have been investigated. PS beads with 750 nm diameter developed a dome-like structure when bubble size varied between 5 μm and 10 μm, beyond which the bubble sizes were too large causing the dome to collapse. In addition, multiple 40 nm AuNPs clusters were simultaneously formed on NPGD array at 16 prescribed locations using a holographic phase SLM. As shown in SEM and LSPR peak mapping, the cluster altered the extinction spectra through plasmonic coupling between AuNP-AuNP and AuNP-NPGD with 50 to 100 nm red shifts compared to the bare NPGD substrate, and 200 nm red shifts compared to non-aggregated, colloidal AuNP. The clusters showed a high SERS active region with intensity amplification 5 times that of bare NPGD which was shown to have a SERS enhancement factor exceeding 10^8^ [31]. We have also demonstrated the feasibility of using this platform to collect and concentrate biological organisms such as microscale *E. coli*. bacteria and nanoscale exosomes.

## Acknowledgement

W.-C.S. acknowledges the National Science Foundation (NSF)CAREER Award CBET-1151154, NSF CBET-1605683, NSF CBET-1643391.

We also acknowledge Dr. Xiaonon Shan and Xu Yang at University of Houston for providing the *E. coli* and Dr. Steven H. Lin at UT MD Anderson Cancer Center for providing the exosome.

